# Saving killer whale populations from a global collapse: rebuttal against Desforges et al. (2018)

**DOI:** 10.1101/474320

**Authors:** Lars Witting

## Abstract

In their *Science* paper, Desforges et al. (2018) address PCB pollution in killer whales, predicting a decline in calf survival and an associated collapse of killer whale populations worldwide. I refute the collapse, showing that it follows from a flawed model parametrisation. The result is not questioning contamination problems in killer whales, but only the bold prediction of global collapse.

---

Desforges et al. (2018) divide killer whales into seven groups from the least PCB polluted in Arctic and Antarctic waters (group 1) to the most contaminated in industrialised regions (group 7), and use a predicted decline in calf survival to forecast killer whale populations over a 100 year period. Population dynamics over 100 years include regulations of the growth rate. Apart from density dependence, the regulation will also adjust the PCB level in individuals as the environmental PCB concentration changes over time.

Being long-lived top-predators, killer whales have a delayed PCB accumulation relative to most animals. It is essential to address whether their current contamination reflects a persistent ecological exposure, or the accumulation of historical pollution in older animals, with subsequent transmission from mothers to calves. Equally important is an environmental model to predict future ecological exposures to PCB. These regulation issues, however, are not addressed by Desforges et al. (2018), who consider unregulated population growth with constant PCB contamination in the different exposure groups. This makes their study unable to predict realistic killer whale dynamics, but it does not imply that they are unable to estimate current impacts on killer whales worldwide.

Their figure 2A illustrates this impact by a calf survival that declines with increased maternal PCB concentration. The response follows from a number of data points, yet these follow from a laboratory response in mink. The predicted population collapse is based on the assumption that we can straightforwardly convert a physiological response in a 0.5 kg terrestrial laboratory animal to natural populations of a free-living marine mammal that is about 10,000 times larger. While it is fair to assume that increased PCB levels will affect calf survival also in killer whales, Desforges et al. (2018) do not address the large degree of uncertainty that is associated with the conversion of a physiological response in mink to a population level response in killer whales.

Assuming that the response in mink is applicable to killer whales, might we, as the authors, conclude that only killer whales *in the less-contaminated waters of the Arctic and Antarctic today appear to be able to sustain growth*. To predict the negative growth of polluted populations, Desforges et al. (2018) use life history data from several populations of killer whales (their Table S2) to develop a base-case of a non-contaminated population. From their predicted 141% increase in abundance over 100 years, we calculate a yearly growth rate of *r* = ln(2.41^1/100^) = 0.88% for their pristine base-case population. This estimate is unexpectedly low, as the growth rate of real killer whale populations—which are exposed to both PCB pollution and density regulation—has been estimated to 2.9% by Olesiuk et al. (1990) and 3.4% by Matkin et al. (2014).

When, as in the case of Desforges et al. (2018), there is no regulation in the projection model, it is essential that the growth of the pristine population relates to some kind of potential, or maximum, estimate. Yet, Desforges et al. (2018) do not attempt to estimate the impacts on potential growth, but only on current growth; which could be zero, or even negative, in a fully viable non-contaminated population. It is flawed to take life history data from a stable natural population and project a collapse by a small downward adjustment of the growth rate in a model with no density regulation. This failure is nevertheless incorporated into the model of Desforges et al. (2018), as some of their life history data are from non-increasing populations of killer whales.

It is equally important to compensate for the detrimental effects of PCB when estimating the pristine base-case from PCB exposed populations. We would otherwise subtract the detrimental effects twice when a smaller than observed calf survival is predicted from the observed PCB level. Desforges et al. (2018) do not apply this correction either when they estimate their non-polluted base-case from PCB polluted populations. The effect from this failure might not be large if the base-case was estimated from the relatively non-polluted Arctic or Antarctic populations. But, it is estimated as an average across nine studies that include medium (Canadian South resident, PCB group 3) and heavily (Strait of Gibraltar, PCB group 7) polluted populations. Their average is furthermore biased towards the medium PCB level of the Canadian South residents, because data from this population—which is non-increasing and composed of only three family groups—are included three times in the average, while data from the much larger and relatively non-polluted populations off Norway, Alaska and the Crozet Archipelago are included only once.

A third issue is the lowest calf survival rate in Table S2—the 0.57 estimate from Olesiuk et al. (1990) for the Canadian residents. This particular calf survival estimate should not be included in the parametrisation because it is absorbed into the fecundity rate (Olesiuk et al. 1990). The hint is that the fecundity rate reflects the number of surviving calves at about 0.5 years of age, while the calf mortality of 0.43 reflects the estimated number of calves that do not survive the first half year, including calves that are dead at birth. Given these issues it is of no surprise that Desforges et al. (2018) estimate a pristine growth rate that is unexpectedly low.

What pristine growth rate should we use to estimate the potential population effects of PCB across the seven contamination groups. A candidate is the 2.9% from Olesiuk et al. (1990), which was estimated for Canadian residents (primarily PCB group 1) before a halt in increase. The underlying rates of increase for separate family groups vary from −0.5% to 4.0%, with the variation originating primarily from differences in the fecundity rate (Brault and Caswell 1993). This type of intra-population variation is expected in natural populations from the density dependent interactive competition between the individuals in the population (Witting 1997, 2017). It implies that the rate of increase in population is approaching the highest intra-population rates when there is minimum interference at low densities. Four percent might thus be a realistic growth potential for killer whales. This is supported by the 4.2% increase that occurred in Alaska killer whales in the 1980s, with the somewhat lower average of 3.4% from 1984 to 2010 being suggested by Matkin et al. (2014) as an estimate of maximum growth.

A range from 2.9% to 3.4% is a realistic, yet somewhat conservative, estimate of potential growth. Figure 1 compares the resulting growth rates across the seven contamination groups, based on the reduced calf survival in Desforges et al. (2018). Where the original study predicted negative growth for contamination groups three to seven, the corrections find potentials for positive growth across the PCB levels of all groups. There is thus no evidence for a global PCB driven collapse of killer whale populations. The observed PCB contamination is nevertheless alarming in light of the increased neonate mortality in polluted minks, and the density regulation of the most exposed killer whale populations may compensate only partly for the detrimental effects of PCB.

**Figure 1:**
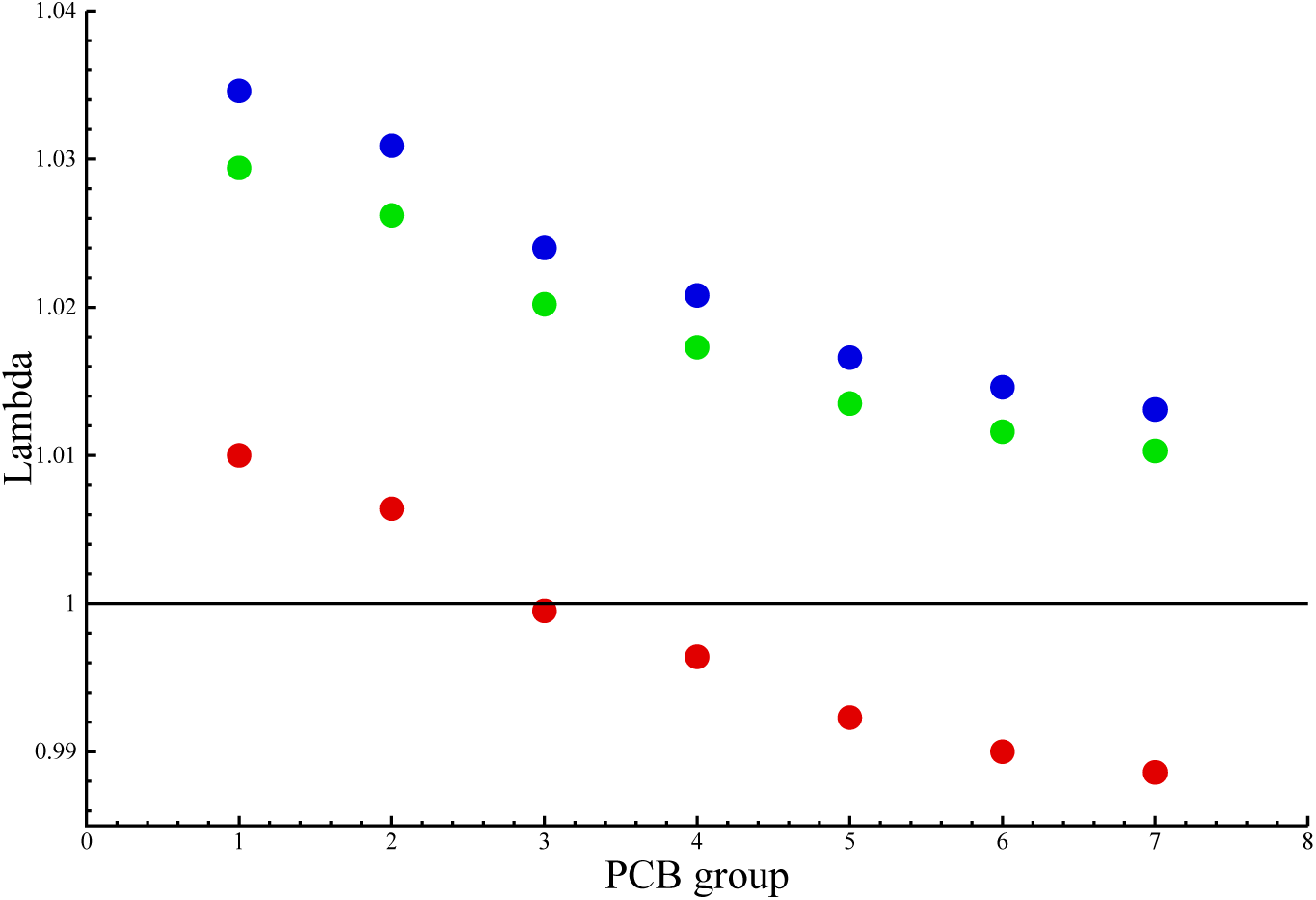
Estimates of population growth (lambda, *λ* = *e^r^*) in killer whale groups exposed to different levels of PCB. Red dots are estimates from Desforges et al. (2018). Blue and green are the corrected estimates when the growth potential of group one is *r* = 3.4% or *r* = 2.9%, as estimated by Matkin et al. (2014) and Olesiuk et al. (1990). The age structured population models are given in the appendix, and the contamination levels of the different groups are defined by Desforges et al. (2018).

## Appendix A

**Table 1:** Life histories that approximate the model in Desforges et al. (2018), following Olesiuk et al. (1990) where calf survival is absorbed in a fecundity rate that declines with increased PCB pollution (G). *p*_0_:Age-class zero survival. *p*_1+_:1+ survival. *a_m_*:Age of first reproduction. *m*:Yearly fecundity. 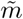:Fecundity relative to group one. *r*:Exponential growth rates from Fig. 2D in Desforges et al. (2018). *m* was solved by the Euler equation to match *r* for the seven contamination groups.

| G | $p_0$ | $p_{1+}$ | $a_m$ | $m$ | $\tilde{m}$ | $r$ |
| --- | --- | --- | --- | --- | --- | --- |
| 1 | 0.951 | 0.966 | 11 | 0.147 | 1.000 | 1.00% |
| 2 | 0.951 | 0.966 | 11 | 0.131 | 0.891 | 0.64% |
| 3 | 0.951 | 0.966 | 11 | 0.104 | 0.706 | -0.05% |
| 4 | 0.951 | 0.966 | 11 | 0.093 | 0.630 | -0.36% |
| 5 | 0.951 | 0.966 | 11 | 0.080 | 0.540 | -0.77% |
| 6 | 0.951 | 0.966 | 11 | 0.073 | 0.493 | -1.00% |
| 7 | 0.951 | 0.966 | 11 | 0.069 | 0.465 | -1.15% |

**Table 2:** The corrected model where *r* = 2.9% and the life history of group one approximate the model in Olesiuk et al. (1990). Relative fecundity (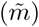) from Table 1.

| G | $p_0$ | $p_{1+}$ | $a_m$ | $m$ | $\tilde{m}$ | $r$ |
| --- | --- | --- | --- | --- | --- | --- |
| 1 | 0.700 | 0.989 | 15 | 0.212 | 1.000 | 2.90% |
| 2 | 0.700 | 0.989 | 15 | 0.189 | 0.891 | 2.59% |
| 3 | 0.700 | 0.989 | 15 | 0.150 | 0.706 | 2.00% |
| 4 | 0.700 | 0.989 | 15 | 0.134 | 0.630 | 1.72% |
| 5 | 0.700 | 0.989 | 15 | 0.114 | 0.540 | 1.34% |
| 6 | 0.700 | 0.989 | 15 | 0.105 | 0.493 | 1.15% |
| 7 | 0.700 | 0.989 | 15 | 0.099 | 0.465 | 1.02% |

**Table 3:** The corrected model where *r* = 3.4% and the life history of group one approximate the model in Matkin et al. (1990). Relative fecundity (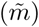) from Table 1.

| G | $p_0$ | $p_{1+}$ | $a_m$ | $m$ | $\tilde{m}$ | $r$ |
| --- | --- | --- | --- | --- | --- | --- |
| 1 | 0.726 | 0.989 | 12 | 0.210 | 1.000 | 3.40% |
| 2 | 0.726 | 0.989 | 12 | 0.187 | 0.891 | 3.04% |
| 3 | 0.726 | 0.989 | 12 | 0.148 | 0.706 | 2.37% |
| 4 | 0.726 | 0.989 | 12 | 0.132 | 0.630 | 2.06% |
| 5 | 0.726 | 0.989 | 12 | 0.113 | 0.540 | 1.65% |
| 6 | 0.726 | 0.989 | 12 | 0.104 | 0.493 | 1.45% |
| 7 | 0.726 | 0.989 | 12 | 0.098 | 0.465 | 1.30% |

## REFERENCES

Brault, S., and H. Caswell 1993. Pod-specific demography of killer whales (Orcinus orca). Ecology 74:1444–1454.

Desforges, J.-P., A. Hall, B. McConnell, A. Rosing-Asvid, J. L. Barber, A. Brownlow, S. De Guise, I. Eullaers, P.D. Jepson, R.J. Letcher, M. Levi, P. S. Ross, F. Samarra, G. Vikingson, C. Sonne and R. Dietz 2018. Predicting global killer whale population collapse from PCB pollution. Science 361:1373–1376.

Matkin, C. O., J. W. Testa, G. M. Ellis and E. L. Saulitis 2014. Life history and population dynamics of southern Alaska resident killer whales (Orcinus orca). Marine Mammal Science 30:460–479.

Olesiuk, P. F., M. A. Bigg and G. M. Ellis 1990. Life history and population dynamics of resident killer whales (Orcinus orca) in the coastal waters of Brithis Columbia and Washington State. Reports of the International Whaling Commission Special Issue 12:209–243.

Witting, L. 1997. A general theory of evolution. By means of selection by density dependent competitive interactions. Peregrine Publisher, Ãrhus, 330 pp, URL http://mrLife.org.

Witting, L. 2017. The natural selection of metabolism and mass selects life-forms from viruses to multicellular animals. Ecology and Evolution 7:9098–9118, http://dx.doi.org/10.1002/ece3.3432.

